## Supplementary material for "Structure of a membrane-bound menaquinol:organohalide oxidoreductase": External Data Material

for

**Extended Data Table 1 -** Cryo-EM data collection, refinement and validation statistics.

| **Data collection and processing** | RDH complex (PceAB) |
|---|---|
| Nominal magnification | 155k x |
| Voltage (kV) | 300 |
| Electron exposure ($e^{-}/Å^{2}$) | 60 |
| Defocus range (-μm) | 1.0-2.2 |
| Physical pixel size (Å) | 0.51 |
| Symmetry imposed | C2 |
| Initial particle images (no.) | 1’049’505 |
| Final particle images (no.) | 34’078 |
| Map resolution (Å) | 2.83 |
| FSC threshold | 0.143 |
| Map resolution range (Å) | 30-2.0 |
| **Refinement** | |
| Initial model used (PDB code) | n/a |
| Map sharpening B factor ($Å^{2}$) | -63.0 |
| Model composition<br>Non-hydrogen atoms<br>Protein residues<br>Ligands | <br>9510<br>1184<br>DCE: 2<br>SF4:4<br>MQ7:2<br>COB:2 |
| R.m.s. deviations<br>Bond lengths (Å)<br>Bond angles (°) | <br>0.004 (4)<br>0.574 (6) |
| Validation<br>MolProbity score<br>Clashscore<br>Poor rotamers (%) | <br>1.32<br>5.82<br>0.00 |
| B-factors (min/max/mean)<br>Protein<br>Ligand | <br>14.07/88.61/35.26<br>22.0/30.78/27.39 |
| Ramachandran plot<br>Favored (%)<br>Allowed (%)<br>Disallowed (%) | <br>98.38<br>1.45<br>0.17 |
| PDB deposition ID | XXX |
| EMDB deposition ID | XXX |
| EMPIAR deposition ID | XXX |

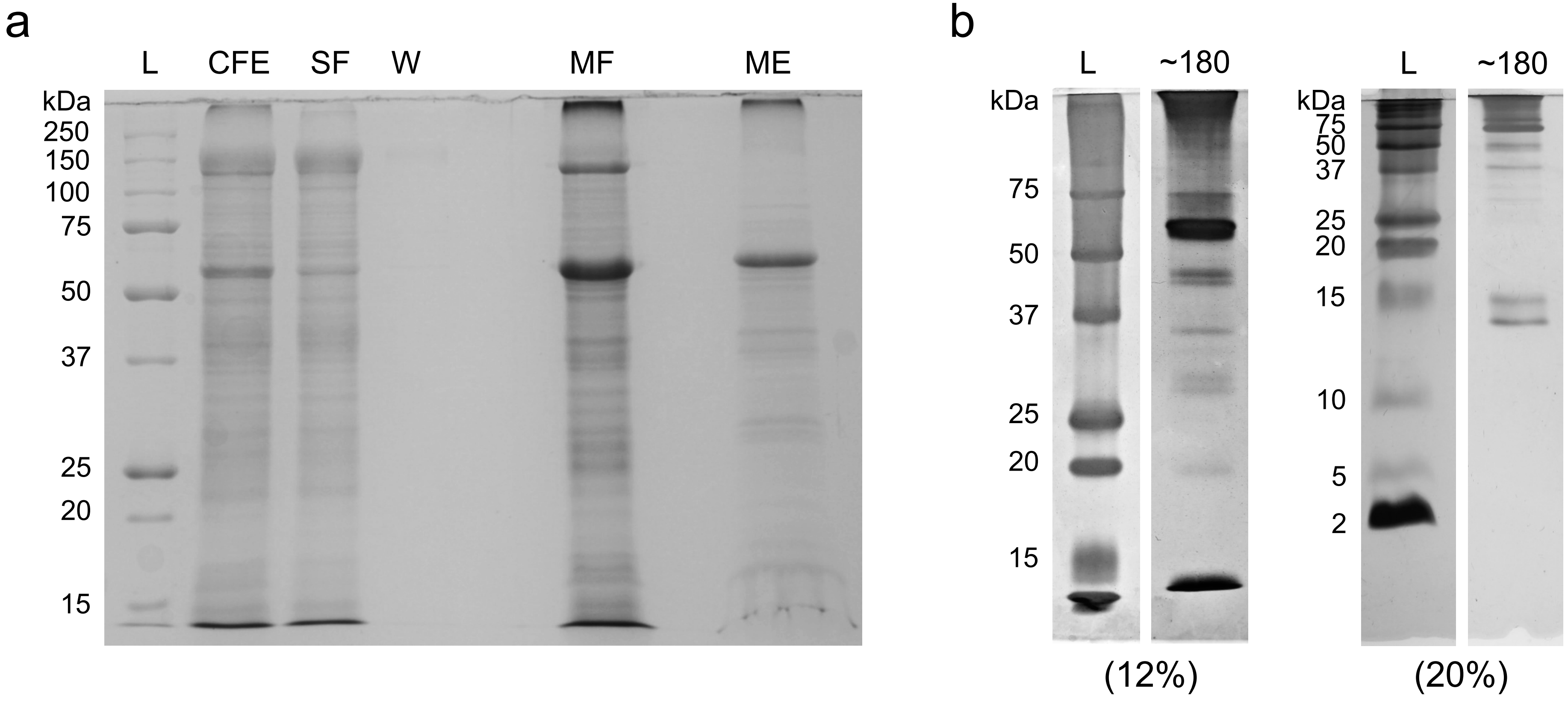


**Extended Data Figure 1 -** SDS-PAGE of **(a)** representative protein samples obtained during the extraction the $PceA_2B_2$ complex from *D. hafniense* strain TCE1, and of **(b)** 2D-electrophoresis of the ~180 kDa band in 12% and 20% acrylamide gels. Legend: L: protein ladder; CFE: cell-free extract; SF: soluble fraction; W: wash sample; MF: membrane fraction; ME: membrane extract.

a

| | Total spectrum count (% coverage) |
|---|---|
| PceA | 2935 (84) |
| PceB | 22 (41) |

b

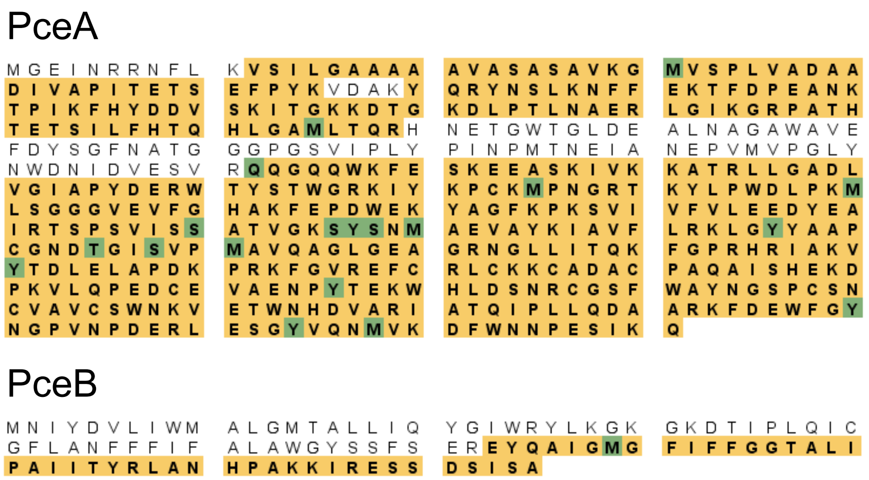


**Extended Data Figure 2 -** LC-MS/MS analysis of the ~180 kDa CN-PAGE band. **(a)** Total spectrum count of PceA and PceB peptides. **(b)** Representative coverage of PceA and PceB proteins highlighted in colour. Green residues indicate those that are possibly modified post-translationally.

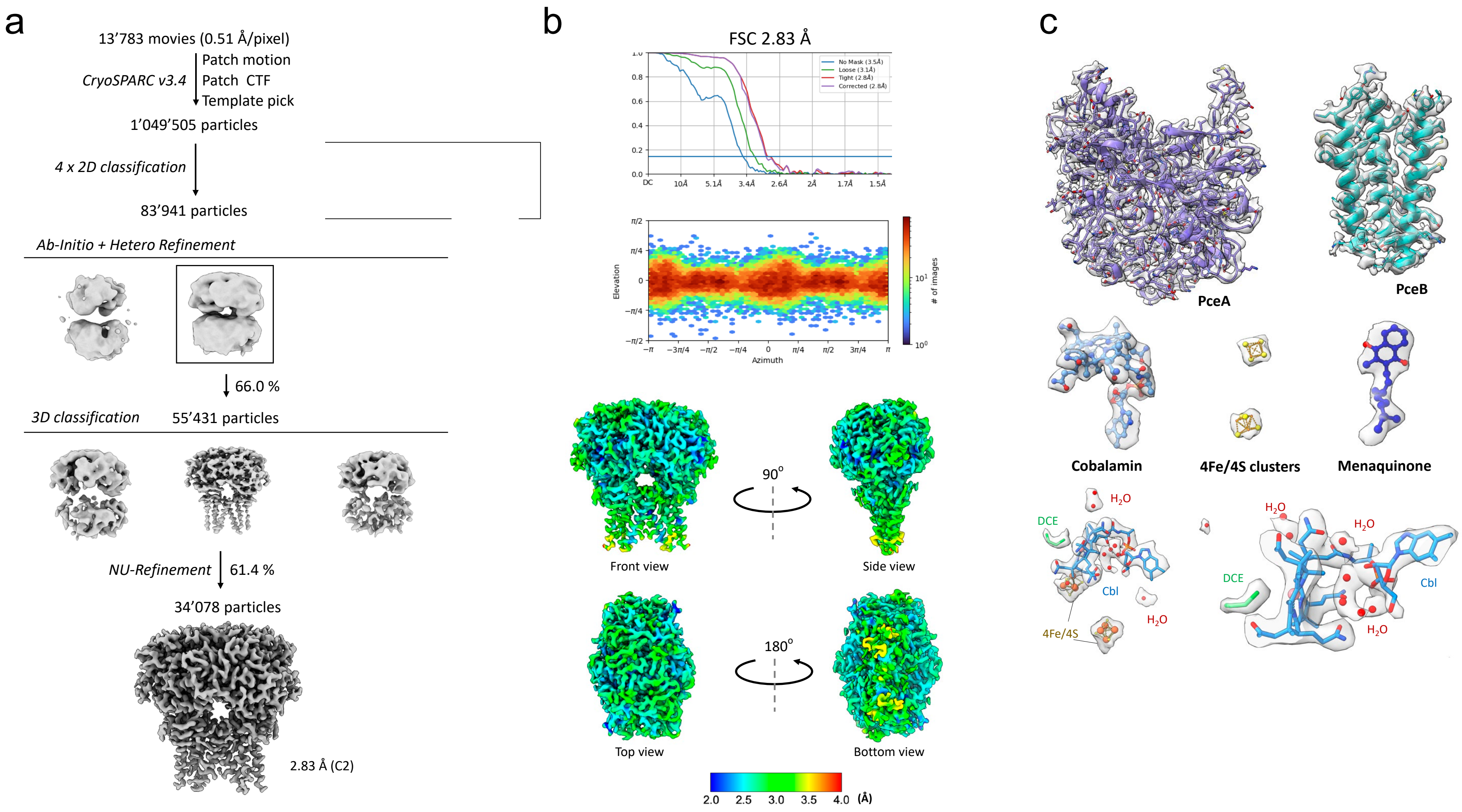


**Extended Data Figure 3** - Cryo-EM analysis of the $PceA_2B_2$ complex. **(a)** Flow chart of the cryo-EM data analysis. **(b)** FSC curve indicating an overall resolution of 2.83 Å (FSC 0.143), direction distribution plot and global refined map coloured by local resolution are given in different views. **(c)** Density maps and structure interpretation of PceA and PceB proteins, and of the cofactors.

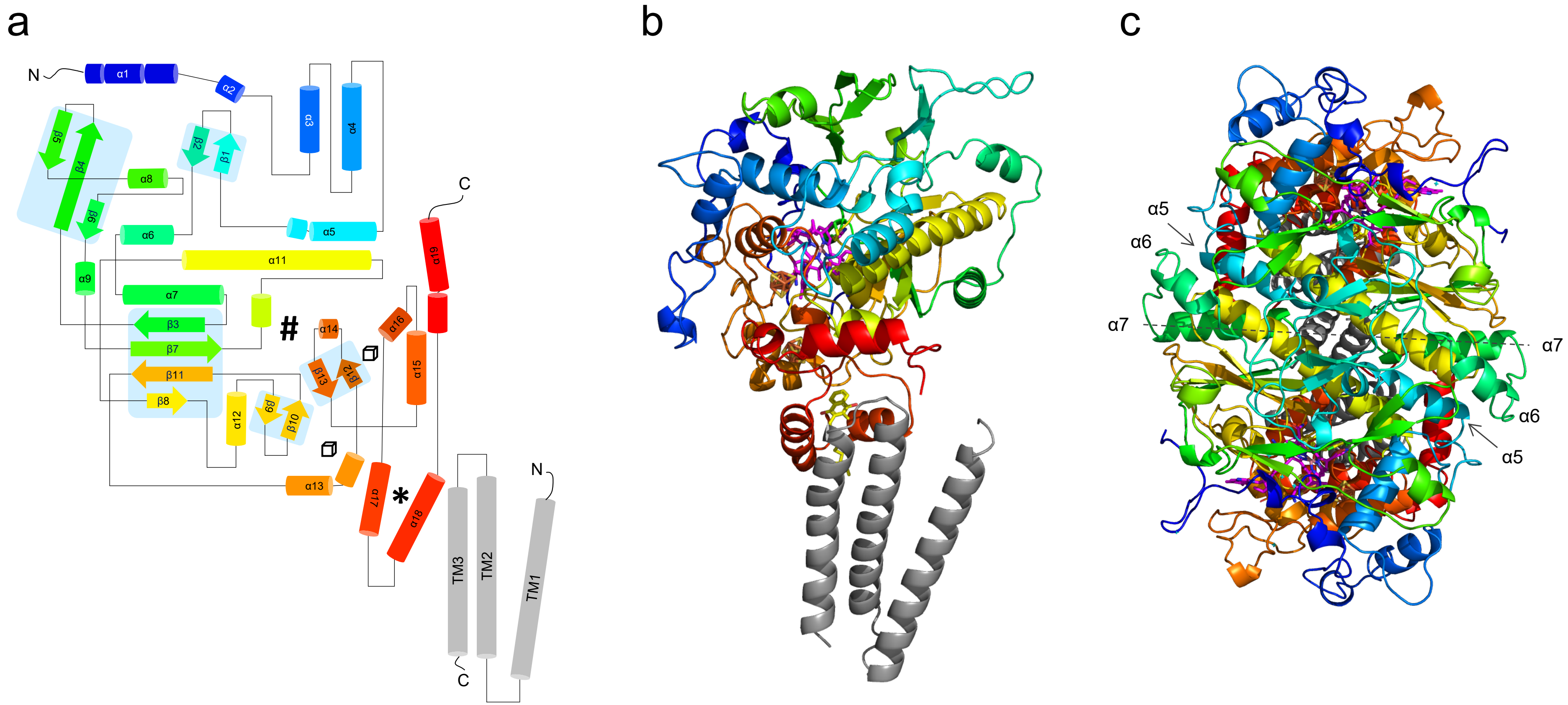


**Extended Data Figure 4 -** Topology analysis of $DhPceA_2B_2$ secondary structures. **(a)** Topology diagram of the DhPceAB heterodimer depicted in rainbow colour mode. **(b)** Front view of the corresponding cartoon structure. **(c)** Top view of the $DhPceA_2$ cartoon structure.

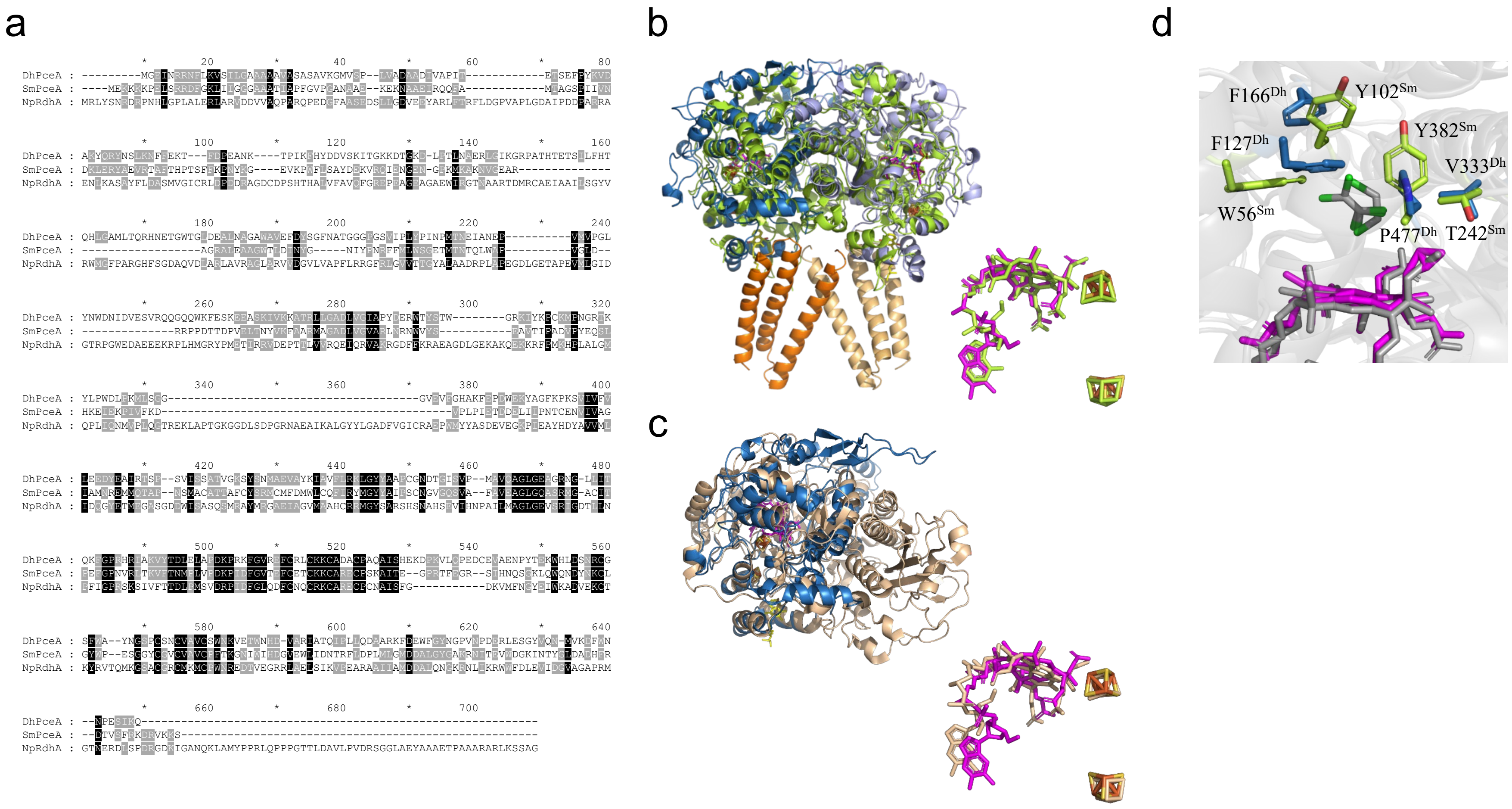


**Extended Data Figure 5** - Sequence and structure alignments of reductive dehalogenases. **(a)** Primary sequence alignment of DhPceA, SmPceA and NpRdhA. Conserved amino acids are shaded in black. **(b)** Structure alignment of $DhPceA_2B_2$ and $SmPceA_2$ depicted in front view. The superimposition of cofactors is also shown. **(c)** Structure alignment of DhPceA and NpRdhA. The cofactors are also shown in a separate panel. **(d)** Active site residue differences between DhPceA and SmPceA. <u>Colour code</u>: $DhPceA_2B_2$ is displayed with the same colours as before, $SmPceA_2$ and its cofactors are in green, while NpRdhA and cofactors in cream.

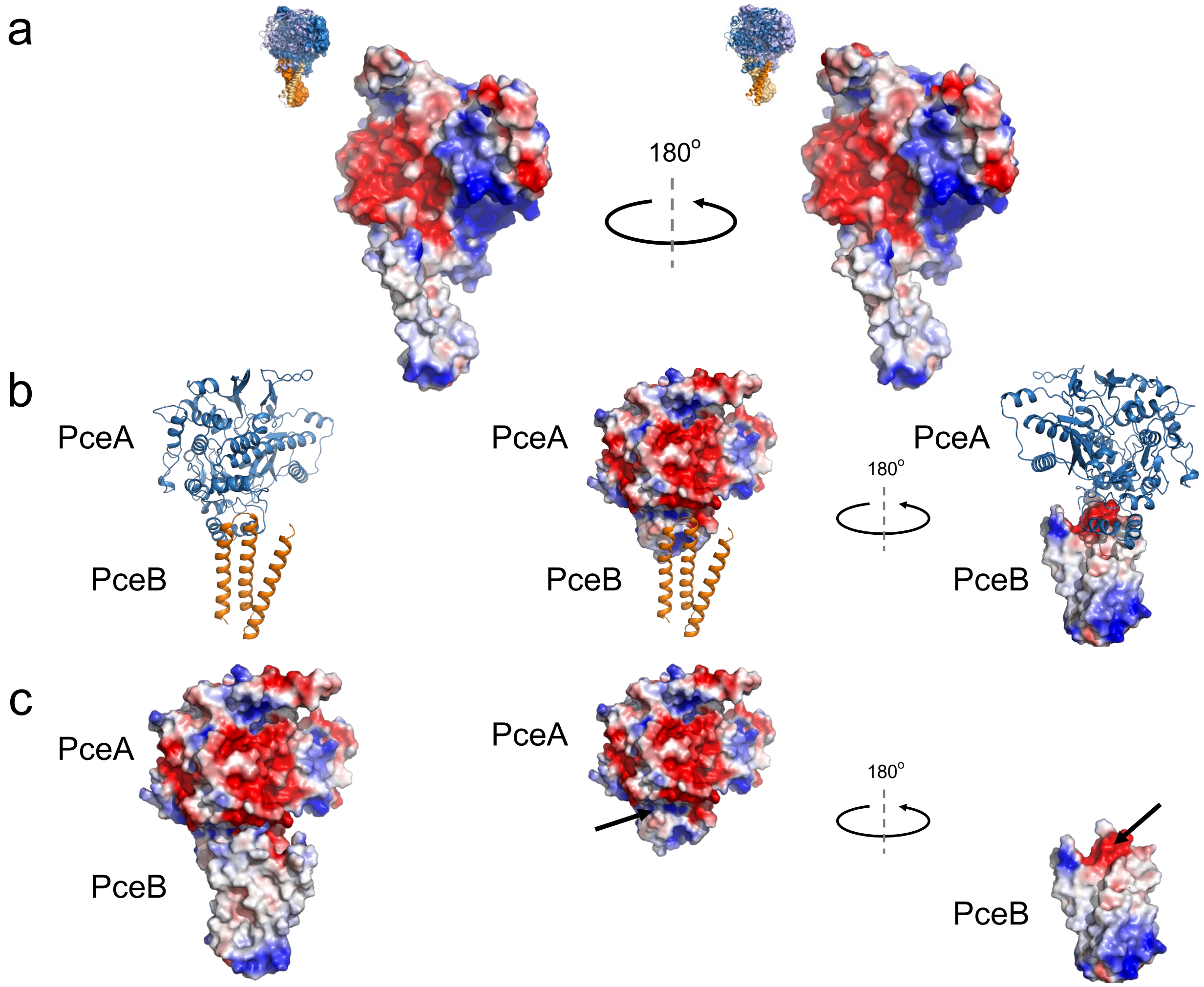


**Extended Data Figure 6 -** Surface interactions within the $PceA_2B_2$ complex. **(a)** Electrostatic interactions between both PceA monomers highlighting the opposite overall charges in the contact area, and hydrophobic interactions between both PceB monomers.
**(b)** and **(c)** Electrostatic interactions between PceA and PceB subunits in each PceAB heterodimer.

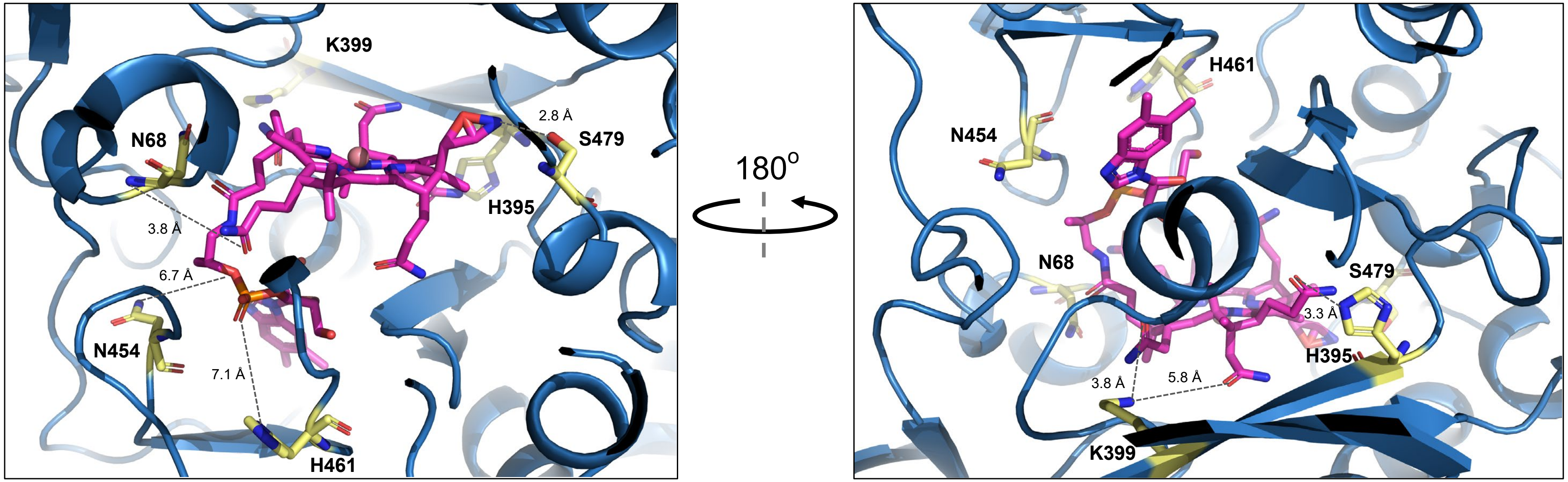


**Extended Data Figure 7 -** Structural insights in the cobalamin binding site of DhPceA showing amino acids at H-bond distance to the cobalamin cofactor.

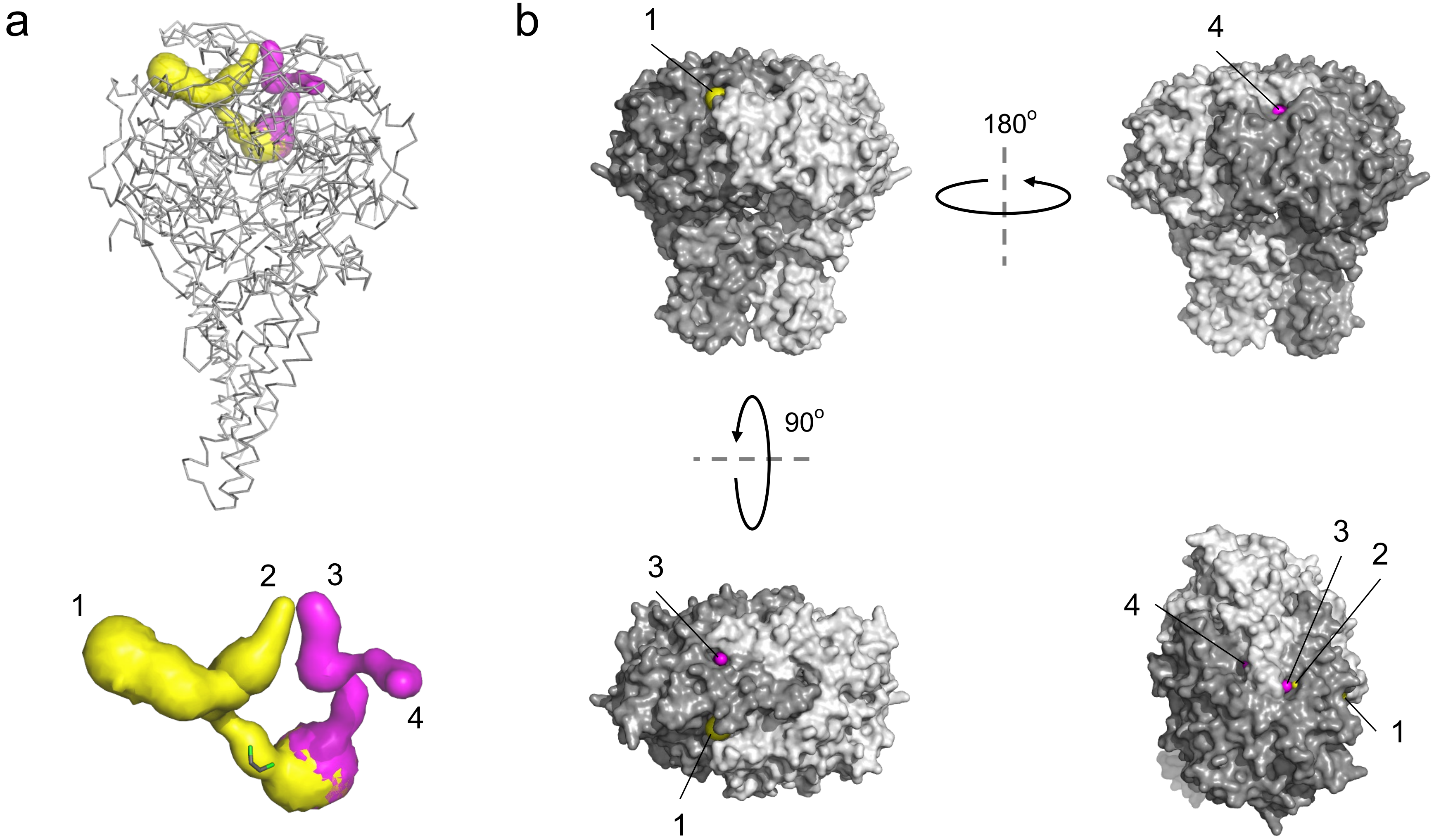


**Extended Data Figure 8 -** Channel prediction in DhPceA. **(a)** Overview of both channels connecting the active site to the outside solution (see also Figure 4 in the article). **(b)** Position of the channel exits at the surface of the DhPce$A_2B_2$ complex. For clarity, the channel exits are only indicated for one PceA subunit (depicted in dark grey).

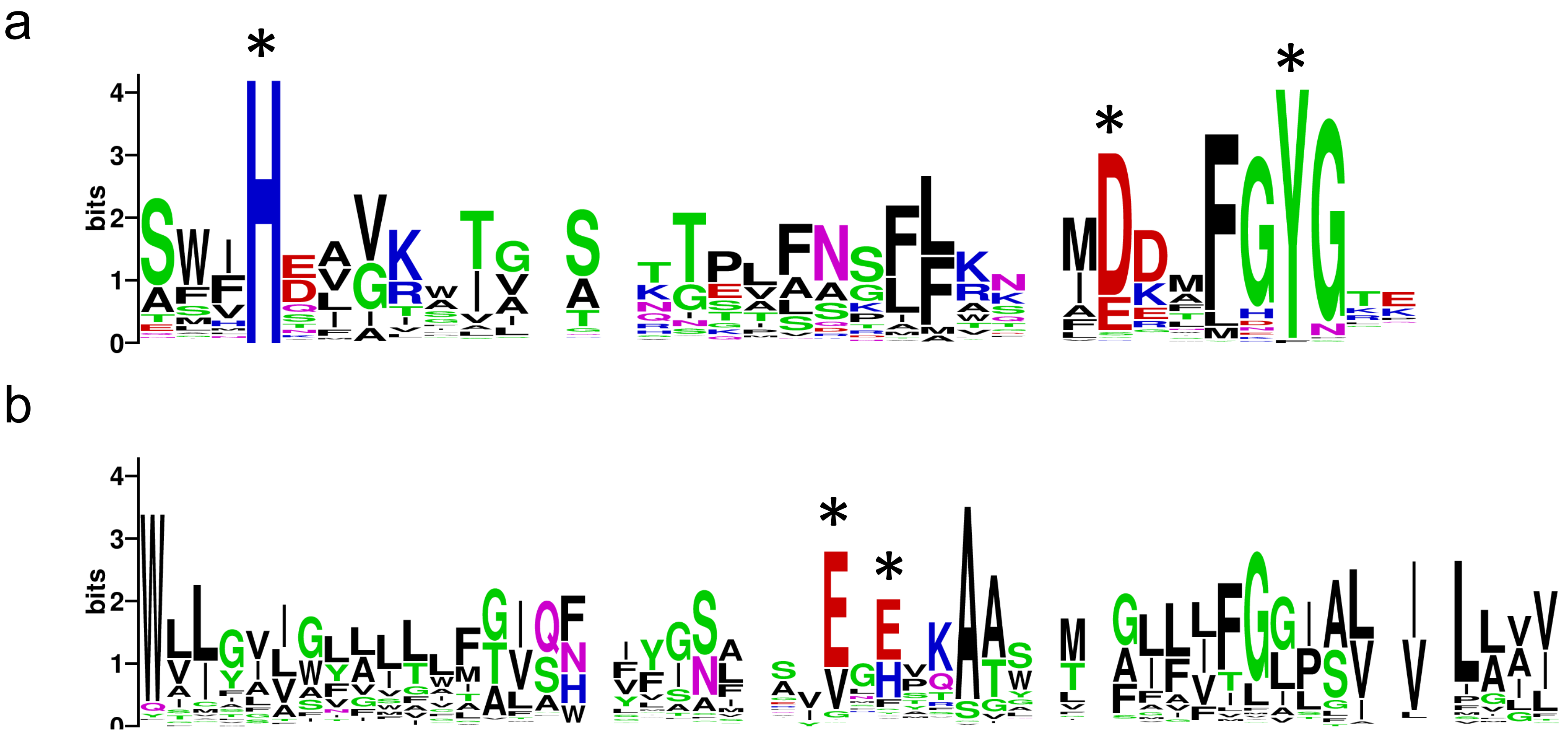


**Extended Data Figure 9 -** Key conserved residues in the MK-binding pocket of RdhAB heterodimers. Weblogo representation of selected regions of **(a)** RdhA and **(b)** RdhB sequence alignments. Stars indicate in the C-terminal region of RdhA sequences a fully conserved histidine residue (H495 in DhPceA) and an almost fully conserved tyrosine (Y520), as well as a highly conserved negatively charged residue (D515). The loop connecting TMH2 and THM3 in RdhB sequences harbours highly conserved glutamate residues (E61 and E63 in DhPceB).
